# Chimeric archaeal BHB-human pre-tRNA scaffold HARBOR system enables versatile production of functional circular RNA modalities

**DOI:** 10.64898/2026.10.12.758436

**Authors:** Shi-Meng Cao, Xin Feng, J.Tian-Chang Zhang, Hao Wu, Xiao Wang, Keqiong Ye, Ling-Ling Chen

## Abstract

Circular RNAs (circRNAs) exhibit superior intrinsic stability and lower immunogenicity compared with linear RNAs, rendering them promising modalities for biomedical and biotechnological applications. Here, we establish the HARBOR (<u>H</u>uman-<u>A</u>rchaeal <u>R</u>NA <u>B</u>ackbone for <u>O</u>verexpression of circular <u>R</u>NA) system, an engineered endogenous circRNA expression platform integrating a human precursor transfer RNA (pre-tRNA) scaffold with an archaeal Bulge-Helix-Bulge (BHB) motif. HARBOR repurposes the endogenous tRNA splicing machinery to generate chimeric pre-tRNA intermediates, which are rapidly degraded following processing, while producing circRNAs ranging from hundreds of nucleotides to 8 knt in cells and *in vivo*. Notably, it enables robust production of large-sized circRNAs with yields nearly 80-fold than the intronic complementary sequence-mediated back-splicing (3D5) system and can drive protein expression from translatable circRNAs at an efficiency close to 20-fold than the Tornado system. Such prominent properties support diverse applications of HARBOR, including producing therapeutic aptamers targeting PKR-associated neuroinflammation and pathological TDP-43 condensation, as well as protein-coding circRNAs encoding adenine base editors for genome editing and MYOD1 for fibroblast-to-myocyte trans-differentiation. Collectively, HARBOR represents a versatile, scalable, and user-friendly platform that unifies circRNA-based aptamer and protein expression applications, advancing circRNA-enabled engineering and biomedical applications.

## Introduction

Circular RNAs (circRNAs) are covalently closed single-stranded transcripts, conferring superior structural stability and inherent resistance to exonuclease degradation^1, 2^. These unique properties make them versatile scaffolds for broad biomedical applications, including programmable protein expression^3, 4^, functional molecular aptamer development^5–9^, and the construction of regulatory RNA circuits^6, 10^. Thus, develop robust and controllable strategies for intracellular circRNA expression is highly desired.

Early circularization strategies in cells mainly relied on two systems: the intronic complementary sequence-mediated back-splicing^2, 11–13^, typified by the 3D5 system^14^, and the system based on archaeal- or metazoan-originated tRNA intron-splicing machineries^15–17^. However, both of them suffer from relatively low circularization efficiency. Subsequent efforts developed the Tornado system, a twister ribozyme-mediated RNA processing platform to efficiently express short circRNA aptamers^6^, and further optimized it to express a translatable circRNA of 4,719 nt using combination of the CMV promoter and CVB3 internal ribosome entry site (IRES)^18^. A robust and simple platform to achieve high-efficiency expression of larger circRNAs remains a challenge. Of note, delivering the *in vitro* transcribed (IVT), ribozyme-driven self-cleaved linear RNAs and employing endogenous cellular RtcB ligase to achieve RNA circularization in theory enables large circRNA expression^19, 20^, however, exogenously delivered linear RNAs may trigger innate immune responses, and the resulting circRNAs may not fully recapitulate the conformation of endogenous ones. Collectively, these challenges highlight the need for a robust circularization strategy rooted in endogenous RNA processing to afford efficient, predictable production of circRNAs harboring large RNA cargos.

During tRNA splicing, the tRNA splicing endonuclease (TSEN) cleaves pre-tRNA at a bulge-helix-bulge (BHB) structural motif. The resulting processing intermediates are then ligated by RtcB to yield a linear mature tRNA and a circular intron^21, 22^. Although the tRNA splicing system has been adapted to generate circRNAs, its efficiency remains low^15–17^. Notably, archaeal pre-rRNAs are also processed by the tRNA splicing machinery into long circular RNAs—exceeding 3,000 nucleotides in the case of pre-23S rRNA^23^. Pre-rRNAs contain highly stable BHB motifs supported by extensive flanking helices^23^, which likely promotes BHB formation and enables efficient long-distance circularization. In contrast, naturally occurring tRNAs typically harbor short introns (<100 nt) and less stable BHB motifs^23^. We hypothesized that incorporating a strong archaeal pre-rRNA BHB motif into tRNA scaffolds may enhance circularization efficiency and expand cargo capacity in tRNA-based circRNA expression platforms.

Here, we developed HARBOR (<u>H</u>uman-<u>A</u>rchaeal <u>R</u>NA <u>B</u>ackbone for <u>O</u>verexpression of circular <u>R</u>NA), a synthetic circRNA expression platform that integrates a human pre-tRNA scaffold with the archaeal pre-rRNA BHB motif. HARBOR supports efficient intracellular production of circRNAs ranging from several hundred to eight thousand nucleotides. We demonstrate that optimized HARBOR system enables *in vivo* generation of functional circRNA aptamers targeting PKR-associated neuroinflammation and pathological TDP-43 condensation. In addition, this platform also empowers protein expression for diverse functional applications, such as adenine base editor (ABE)-mediated genome editing and MYOD1-driven fibroblast-to-myocyte trans-differentiation. Collectively, HARBOR is a robust and versatile platform for functional circRNA expression, expanding the current toolkit for RNA-based synthetic biology and therapeutic development.

## Results

### Most natural tRNA scaffolds perform poorly in circRNA production in cells

To profile the capacity of BHB motifs to mediate circRNA production in mammalian cells, we selected a panel of such motifs from diverse species and RNA classes, and tested their capacity to circularize *circPOLR2A(9,10),* a double-stranded region-containing circRNA (ds-cRNA) known to inhibit PKR hyperactivation^5, 7–9^. Of note, the tested motifs comprised two archaeal pre-tRNA-derived BHB motifs, two archaeal pre-rRNA-derived BHB motifs, and two human pre-tRNA-derived BHB-like structural elements (Fig. 1a). All archaeal BHB motifs and human pre-tRNA structures were symmetrically inserted flanking the linearized *circPOLR2A(9,10)* sequence to generate a panel of circRNA expression constructs.

**Fig. 1.**
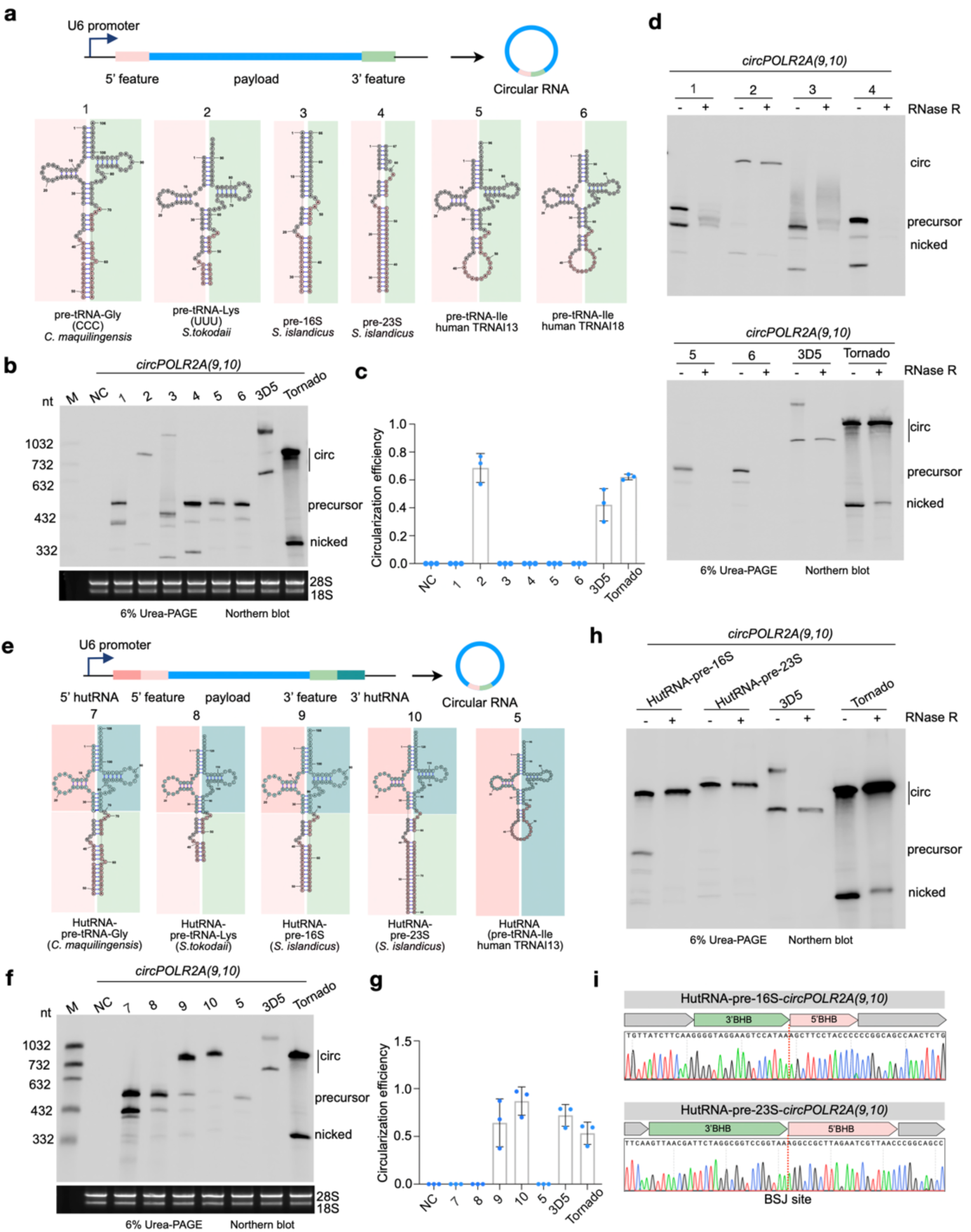
Human pre-tRNA scaffold combined with archaeal BHB motif enables efficient circular RNA formation. **a**, Schematic illustration of the plasmid construction strategy and the structural features of BHB motifs from different origins and various tRNAs. The colored sequence blocks on both sides represent flanking regions of the payload. **b**, Circular RNA formation mediated by each plasmid is shown in panel **a**, as well as by the 3D5^14^ and Tornado^6^ systems, shown by 6 % denaturing PAGE Northern blotting. M, marker; NC, negative control plasmid. 3D5 and Tornado as positive control plasmids. “circ” is RNA circle; “precursor” is linear precursor; “nicked” is nicked circle. **c**, Quantification of circular RNA circularization efficiency mediated by different plasmids shown in panel **a** and panel **b**. **d**, Among all six vectors, only the construct 2 enables an efficient circularization. RNA circular bands were validated by RNase R treatment and indicated by the blue circle. **e**, Schematic of the plasmid construction strategy combining a human tRNA scaffold with individual archaeal BHB motifs. **f**, Northern blotting of circular RNA products mediated by plasmid constructs shown in panel **e**, as well as by 3D5 and Tornado systems. **g**, Quantification of circularization efficiency mediated by individual vectors shown in panel **f**. **h**, Combination of BHB motifs derived from *S. islandicus* pre-16S and pre-23S rRNA with a human tRNA scaffold enables efficient RNA circularization. The formation of circular RNA was validated by RNase R treatment followed by northern blotting assay. **i**, Sanger sequencing confirms the back-splice junction of circular RNA generated by combining a human tRNA scaffold with BHB motifs derived from *S. islandicus* pre-16S and pre-23S rRNA. In panel **c** and **g**, error bars represent the mean ± s.d.

Following transfection of individual chimeric construction into HeLa cells, Northern blot analysis was performed to examine *circPOLAR2A(9,10)* formation. The archaeal pre-rRNA BHB motifs by themselves do not support splicing in cells, consistent with human TSEN recognizing not only BHB motif but also the entire structure of pre-tRNA^24–26^. Among all pre-tRNA candidates tested, only the BHB motif derived from the *Sulfolobus tokodaii* (*S. tokodaii*) pre-tRNA efficiently drove RNA circularization, whereas no circRNA formation was observed from the remaining candidates (Fig. 1b). Compared to the reported 3D5^14^ and Tornado system^6, 18^, the *S. tokodaii* BHB motif achieved a comparable circularization efficiency (circRNA signal relative to total RNA signal), although the overall circRNA yield was markedly lower (Fig. 1b, 1c). Further RNase R digestion showed the exonuclease resistance of circularized products, confirming their closed circular topology (Fig. 1d). As most natural tRNAs perform poorly in expressing circRNA, we therefore sought to further engineer tRNA scaffold to improve circularization efficiency and yield.

### A human tRNA scaffold with archaeal pre-rRNA BHB motif enhances circRNA production

To optimize circRNA expression efficiency in mammalian cells, we hypothesized that combining a human tRNA scaffold with an archaeal BHB motif might synergistically enhance circularization. We selected the precursor transcript of the human tRNA-Ile gene *TRNAI13*^27^ as the core scaffold and incorporated four archaeal BHB motifs derived from archaeal pre-tRNAs or pre-rRNAs to generate a panel of chimeric constructs. The native *TRNAI13* precursor (HutRNA) was served as a control (Fig. 1e).

As shown by Northern blotting, combining the human pre-tRNA scaffold with individual archaeal 16S or 23S pre-rRNA-derived BHB motif markedly improved circRNA formation (Lanes 9, 10) relative to the HutRNA alone control (Lane 5) (Fig. 1f, 1g). In contrast, incorporation of archaeal pre-tRNA-derived BHB motifs into the same human pre-tRNA scaffold failed to produce detectable circularization products (Lanes 7, 8) (Fig. 1f, 1g). RNase R treatment confirmed the circular identity of the RNA products with reduced precursors and nicked bands, while the circRNA levels have remained unchanged (Fig. 1h). Sanger sequencing further confirmed the precise back-splice junction (BSJ) site in each circRNA generated from 16S and 23S pre-rRNA BHB-containing constructs (Fig. 1i). These results strongly suggest that embedding certain archaeal rRNA-derived BHB motif into a human tRNA scaffold enhances engineered circRNA biogenesis in cells.

### BHB optimization for compact and efficient circRNA producing constructs

The original archaeal 23S rRNA-derived BHB motif introduced 47 nt of exogenous sequence into mature circRNAs (Fig. 1i). To minimize exogenous sequence insertion without sacrificing high circularization activity, we designed seven truncated variants targeting this extended duplex region. Each variant featured progressive truncation of the duplex by 2 base pairs (bp), achieved via symmetric deletion of 2nt from both ends of the stem (Fig. 2a). Among all constructs, only the 23S-m1 variant carrying a slightly reduced 43 nt sequence exhibited the high circularization efficiency, even surpassing the wild-type BHB motif. Further assays showed that although the original design introduced 47 nt of exogenous sequence, at least 31 nt of the 23S rRNA-derived insert is required for circularization (Fig. 2b, 2c).

**Fig. 2.**
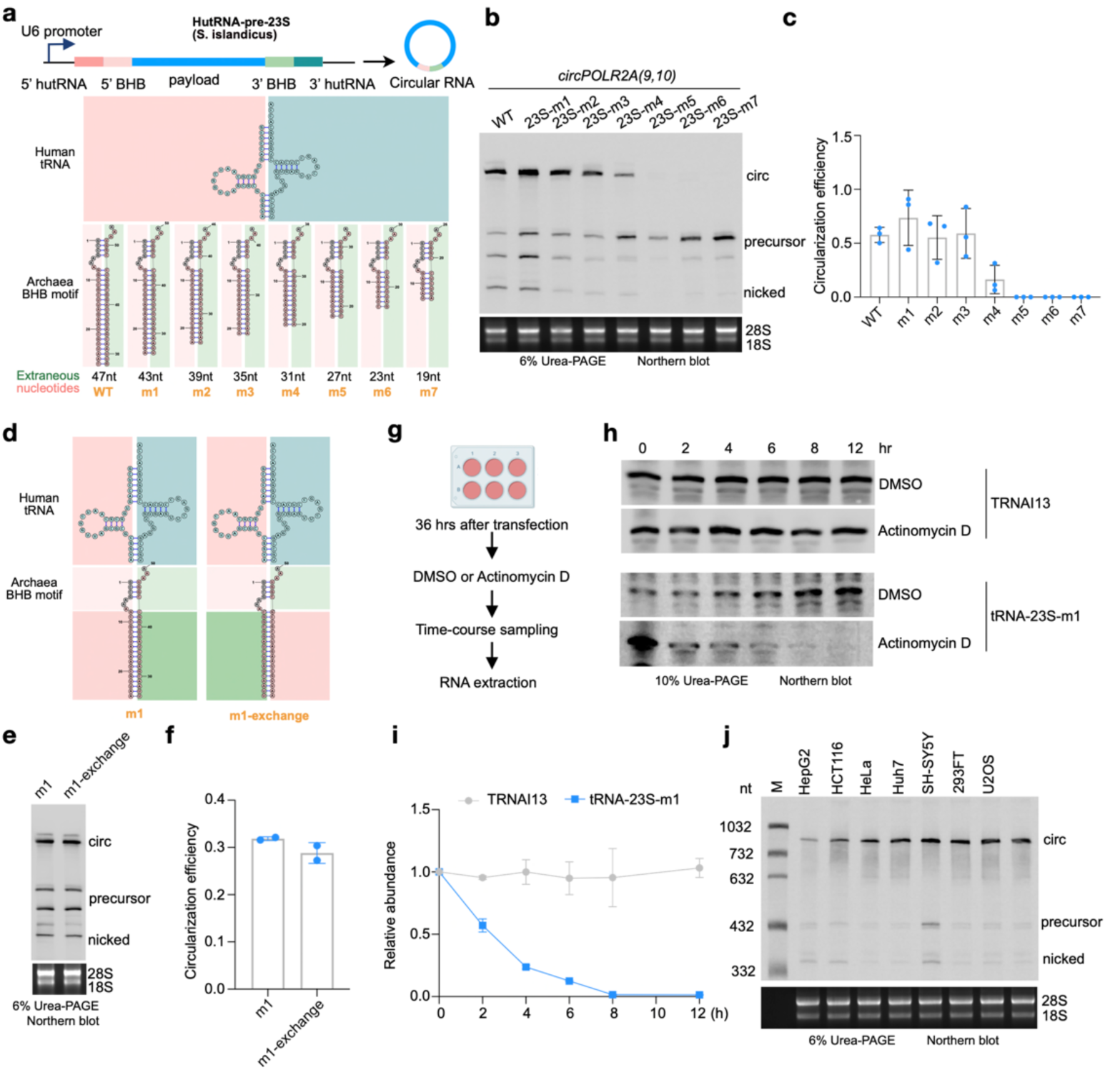
Optimization and characterization of the HARBOR system. **a**, Schematic of truncation mutations in the hutRNA-pre-23S plasmid construct, in which the RNA paring region is progressively shortened by four nucleotides and designated as m1, m2, m3, m4, m5, m6, and m7, respectively. **b**, Northern blotting of circular RNA products mediated by truncation mutants shown in panel **a**. **c**, Quantification of circularization efficiency mediated by individual plasmid vectors shown in panel **b**. **d-f**, Circular RNA formation depends on the structural configuration of the BHB motif rather than its sequence. Of note, reciprocal exchange of the sequences flanking the RNA pairing region does not affect circularization efficiency. **D**, Schematic of the specific construct modifications. **E**, Northern blotting to show circular RNA formation by each indicated construct in cells. **f**, Quantification of the circularization efficiency. **g-i**, HARBOR-derived tRNA intermediates are rapidly degraded. **g**. Schematic illustration of the experimental workflow for measuring the half-life of HARBOR-derived tRNA intermediates. **h**, Endogenous tRNA and HARBOR-produced tRNA (tRNA-23S-m1) were analyzed by northern blotting at indicated time points to assess RNA half-life. Cells treated with DMSO served as the control. Quantification of tRNA half-life is shown in panel **i**. **j**, HARBOR enables efficient circular RNA formation across cell lines. HARBOR plasmids were transfected into indicated cell lines, followed by RNA extraction and analysis by Northern blot to examine circularization efficiency. In panel **c, f** and **i**, error bars represent the mean ± s.d.

In parallel, the archaeal 16S rRNA-derived BHB motif naturally introduced only 31 nt of exogenous sequence. We asked whether this region could be further shortened without compromising circularization capability. We generated truncation mutants by stepwise truncating the extended duplex outside the predicted catalytic core with the same strategy as above (Fig. 2a, Extended Data Fig. 1a). Northern blot quantification showed that all truncated 16S variants displayed reduced circularization efficiency compared to wild type, with the circularization progressively declining as truncation increased (Extended Data Fig. 1b, 1c), highlighting the importance of the intact BHB motif for circRNA engineering.

These results together indicate that the extended duplex region is indispensable for efficient circRNA production, likely by stabilizing the higher-order BHB architecture required for proper RNA processing. Given that the 23S-m1 construct exhibited the highest circularization efficiency (Fig. 2a–2c), we selected this construct for following studies and referred it as HARBOR (<u>H</u>uman-<u>A</u>rchaeal <u>R</u>NA <u>B</u>ackbone for <u>O</u>verexpression of circ<u>R</u>NA).

### HARBOR expresses circRNAs in a structure-dependent manner

To dissect whether the extended duplex region governs circularization via primary sequence or secondary structure, we exchanged the complementary sequences within the duplex while maintaining overall stem-loop architecture (Fig. 2d). Northern blot assays confirmed that sequence-swapped constructs retained circularization efficiency equivalent to the original scaffold (Fig. 2e, 2f), suggesting that BHB-mediated circRNA formation is largely independent of primary sequence and instead depends on duplex architecture. This is consistent with BHB-mediated processing, where substrate recognition is primarily determined by RNA secondary structure rather than sequence identity.

Endogenous pre-tRNA splicing is executed by the TSEN endonuclease complex, which excises intronic segments to generate exon halves that are subsequently ligated by the RtcB ligase to form mature tRNAs^21^. HARBOR repurposes this endogenous splicing machinery, which also generates chimeric tRNA intermediates composed of the human tRNA scaffold and a synthetic archaeal BHB-derived anticodon stem-loop. We then asked whether such engineered chimeric tRNAs was recognized by cellular RNA surveillance pathways. To test this view, we monitored the stability of HARBOR-derived tRNA intermediates (tRNA-23S-m1) upon transcriptional inhibition with actinomycin D in HeLa cells (Fig. 2g), with DMSO-treated cells serving as controls. The results showed that the endogenous *TRNAI13* tRNA remained highly stable over the time course, whereas HARBOR-produced chimeric tRNA intermediates underwent rapid degradation with half-life of ∼2 h (Fig. 2h–2i). These results suggest that HARBOR produced tRNA intermediates do not accumulate thus unlikely perturb endogenous translational homeostasis.

Finally, we evaluated the compatibility of HARBOR across a panel of cell lines, including HepG2, HCT116, HeLa, Huh7, SH-SY5Y, 293FT, U2OS and A549 cells. Northern blotting confirmed robust circRNA expression in all tested cell types (Fig. 2j), indicating HARBOR as a robust circRNA production platform in mammalian cells.

### HARBOR enables high-capacity production of large circRNA cargoes

To define the cargo capacity of HARBOR, we tested a panel of RNAs ranging from 336 nt to 8,000 nt, including *circPOLR2A(9,10)* (336 nt), *circ-sgArray* (452 nt), *circ-CVB3-FLAG* (771 nt), *circ-CVB3-mCherry* (1,476 nt)*, circRTN4(2,3)* (2,457 nt), *circ-CVB3-PpLuc* (2,463 nt), *circ-SV-A-saKKH-ABE8e* (4,632 nt), *circ-CVB3-ABE8e* (5,553 nt), *circ-SV-FHB-DMD* (6,748 nt) and *circ-DMD-8K* (8,000 nt) (Fig. 3a). Northern blotting combined the BSJ site sequencing confirmed the accurate and efficient circularization across all constructs (Fig. 3b, Extended Data Fig. 2a). Notably, the fact that HARBOR efficiently generates circRNAs up to 8,000 nt of our tested cargos, outperforming all described circRNA expression platforms in cells^6, 18^. These findings together show HARBOR as a robust system for delivering different circRNA payloads with distinct purposes.

**Fig. 3.**
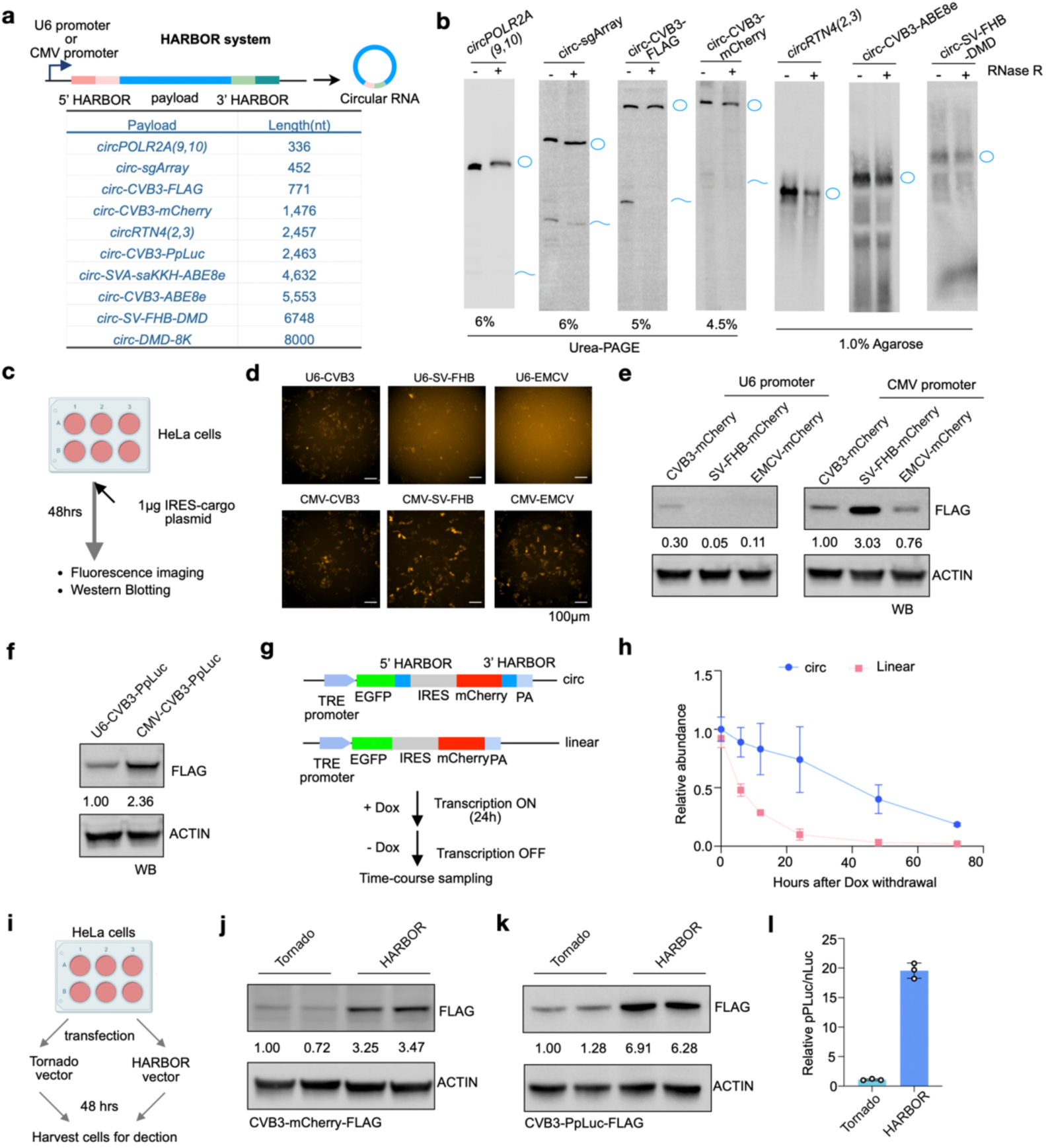
HARBOR vector enables circularization and expression of RNA up to 8 knt. **a**, Schematic of circular RNA cargos with different lengths expressed by the HARBOR system. **b**, Northern blotting of circular RNA products generated by HARBOR. Circular RNAs were validated by RNase R treatment and indicated by blue circles. **c-e**, HARBOR enables an efficient protein expression. **c**, Schematic of HeLa cells transfected with the HARBOR vector for expression of mCherry protein. **d**, mCherry expression driven by different promoters and IRES elements shown by Fluorescence microscopy images; scale bar, 100 μm. **e**, Western blotting of mCherry protein expression under different promoter and IRES configurations. **f**, Western blotting of PpLuc luciferase expression under different promoter configurations. **g-h**, HARBOR-derived circular RNAs exhibit higher stability than linear RNAs generated from mutant constructs lacking the 5′ and 3′ terminal HARBOR sequences. **g**, Schematic of circular RNA and linear RNA expression constructs. Cells were transiently transfected with the two plasmids, followed by doxycycline (dox) induction for 24 h to initiate transcription; dox was then withdrawn and cells were harvested at the indicated time points. **h**, Comparison of RNA half-life between circular RNA and linear RNA produced from individual constructs shown in **g**. **i-l**, HARBOR mediates higher protein expression than the Tornado system ^6^. **i**, Cells were transfected with individual HARBOR- or Tornado-based constructs, followed by protein expression analysis. **j-k**, mCherry (**j**) and PpLuc luciferase (**k**) expression is higher by HARBOR than Tornado, as shown by Western blotting. **l**, The luciferase activity of the PpLuc reporter is higher in HARBOR-transfected cells than in Tornado-transfected cells, as measured by luminescence assay. In panel **h** and **i**, error bars represent the mean ± s.d.

### HARBOR supports efficient cap-independent protein expression from circRNAs

Given the efficient production of large-sized circRNAs from HARBOR, we next characterized the translational competence of such HARBOR-circRNAs. We generated HARBOR constructs encoding mCherry with two different promoters (U6 or CMV), combined them with three distinct IRESs (CVB3, SV-FHB, or EMCV)^4^, and evaluated their translation efficiency in HeLa cells (Fig. 3c).

Fluorescence imaging and immunoblotting showed that CMV-driven constructs yielded higher mCherry protein output than U6-driven counterparts (Fig. 3d, 3e), and a comparable trend was also observed for HARBOR-driven *Photinus pyralis* luciferase (PpLuc) expression (Fig. 3f). Among the promoter-IRES combinations tested, the CMV-SV-FHB configuration yielded the highest protein output (Fig. 3d, 3e). Intriguingly, short circRNA cargos such as *circPOLR2A(9,10)* exhibited markedly lower expression under the CMV promoter than the U6 promoter (Extended Data Fig. 2b). We hypothesized that it may be attributed to the robust transcription driven by CMV that perturbed co-transcriptional RNA folding, thereby impairing TSEN-mediated RNA processing. Together, these results suggest that HARBOR enables both coding and noncoding circRNA expression in mammalian cells, with tunable output via rational promoter and IRES design.

We further compared RNA stability between HARBOR-derived circRNAs and their linear counterparts using a Tet-On inducible system (Fig. 3g). Following 24 h doxycycline induction, transcription was shut off, and RNA decay kinetics was monitored for over 72 h. We observed that more than 70% of HARBOR-derived circRNAs remained detectable 24 h after transcriptional shutoff, whereas the corresponding linear transcripts rapidly decayed to approximately 10% of their initial levels over the same period (Fig. 3h). These results demonstrate the markedly enhanced stability of HARBOR-derived circRNAs, consistent with the intrinsic resistance of circular RNA topology to exonuclease-mediated degradation.

### HARBOR outperforms existing circRNA expression platforms

We next compared HARBOR with the established 3D5^14^ and Tornado^6^ systems for circRNA expression in cells. As shown above (Fig. 1f), HARBOR displayed robust expression of short circRNAs, such as *circPOLR2A(9,10)*, with performance comparable to Tornado, but remarkably outperformed the 3D5 system. We then evaluated their capacity to express large circRNAs. Tornado system has been reported to generate a translatable circRNA of 4,719 nt^18^, while HARBOR supported circularization of transcripts as large as 8 knt (Fig. 3a, 3b; Extended Data Fig. 2a), showing its superior large-cargo capacity.

Given that long circRNAs are frequently used for protein expression, we compared the translational performance of HARBOR and Tornado constructs. HARBOR- and Tornado-based expression plasmids encoding CVB3-mCherry or PpLuc were generated and transfected into HeLa cells, followed by quantification of protein expression levels (Fig. 3i). HARBOR-mediated constructs consistently outperformed the corresponding Tornado systems, producing much higher levels of both mCherry and PpLuc proteins (Fig. 3j, 3k). Notably, PpLuc reporter activity was increased by close to 20-fold in the HARBOR system relative to Tornado (Fig. 3l), highlighting its superior translational performance and utility as a robust circRNA expression platform (Extended Data Fig. 2c).

### HARBOR-derived aptamer exerts neuroprotective effects in AD models

Leveraging the robust intracellular circRNA expression of HARBOR, we next explored its capability to express therapeutic circRNA aptamer *in vivo*. Our previous study identified that 3D5-derived *circPOLR2A(9,10)* efficiently suppressed aberrant PKR activation, thereby mitigating the neuroinflammation and neurotoxic protein accumulation, ultimately improving learning and cognitive performance in AD mouse models^9^. We first evaluated whether HARBOR-derived *circPOLR2A(9,10)* could inhibit aberrant PKR activation in cultured cells (Fig. 4a, Extended Data Fig. 3a, 3b). A549 cells were transfected individually with constructs expressing 3D5-circ*POLR2A(9,10)*^9^ and HARBOR-*circPOLR2A(9,10)*. 24h post transfection, cells were stimulated with poly(I:C) for 2 h to mimic inflammatory stress followed by PKR activation detection (Extended Data Fig. 3a). Consistent with its higher expression level, HARBOR-derived *circPOLR2A(9,10)* suppressed PKR hyperactivation more effectively than the corresponding 3D5-derived circRNA (Extended Data Fig. 3b).

**Fig. 4.**
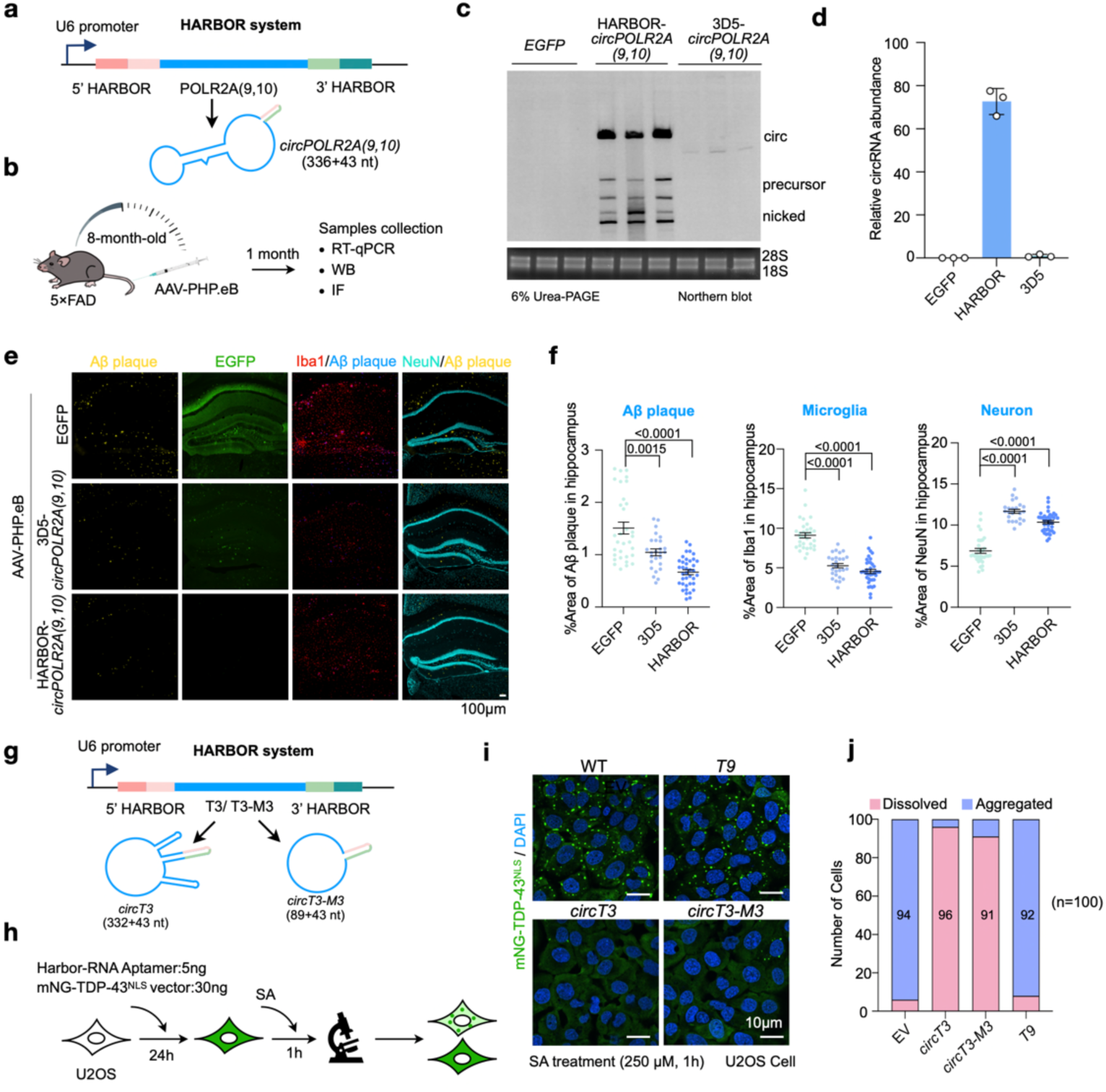
HARBOR displays high efficacy to mitigate circRNA aptamer-targeted pathogenic proteins. **a**, Schematic of HARBOR-mediated *circPOLR2A(9,10)* expression. **b**, Schematic of the experimental design. AAV-PHP.eB-delivered HARBOR-*circPOLR2A(9,10)* was injected into the hippocampus of 8-month-old 5×FAD mice. Samples were collected one month after AAV injection for analysis as previously described^9^. **c**-**d**, AAV-HARBOR produces approximately 80-fold higher circRNA levels than AAV-3D5, as revealed by Northern blotting (**c**) and quantified in (**d**). Error bars represent the mean ± s.d. **e-f**, Reduced neuronal loss, alleviated gliosis, and decreased Aβ accumulation in the hippocampus of 8-month-old 5**×**FAD mice, one month after AAV-delivered HARBOR-*circPOLR2A(9,10)* or AAV-3D5-*circPOLR2A(9,10)* injection. Of note, HARBOR-*circPOLR2A(9,10)* shows a stronger effect than the previously reported AAV-3D5-*circPOLR2A(9,10)* system^9^. **e**, Representative images of Aβ plaques (yellow), EGFP (green), Iba1 (red), and NeuN-labeled neurons (cyan) in hippocampus 5**×**FAD mice with above treatments. Scale bar, 100 μm. **f**, Quantification showing reduced neuronal loss, decreased microglial activation, and reduced Aβ accumulation compared with the EGFP control group. Of note, the AAV-HARBOR-*circPOLR2A(9,10)* group showed better efficacy of all examined parameters than the AAV-3D5-*circPOLR2A(9,10)*-treated group. In panel **f**, statistical analysis was performed using a two-tailed Student’s t-test. Error bars represent the mean ± s.e.m. **g**, Schematic of HARBOR-mediated *circT3* and *circT3-M3* expression. **h**, Schematic of the use of HARBOR-expressed RNA aptamers to prevent TDP-43 cytosolic condensation in U2OS cells. **i**, Representative images of U2OS cells expressing HARBOR-delivered RNAs (*T9*, *circT3*, and *circT3-M3*) or empty vector (EV) as a control, showing their inhibitory effects on TDP-43 condensation. mNG-TDP-43 is shown in green; nuclei are stained with DAPI. Scale bar, 10 μm. **j**, The statistics of TDP-43 condensation from panel **h**, showing that *circT3* and *circT3-M3* RNAs effectively inhibit cytosolic aggregation.

Next, we compared the *in vivo* expression efficiency and therapeutic efficacy of 3D5 and HARBOR systems. *circPOLR2A(9,10)* expressed by these two systems were individually delivered to 8-month-old 5×FAD AD mice^9, 28^ via AAV-PHP.eB administration (Fig. 4b). Notably, HARBOR-driven *circPOLR2A* achieved robust i*n vivo* expression, exceeding that of the 3D5 system by close to 80-fold (Fig. 4c, 4d). To minimize potential dose-dependent safety concerns associated with AAV administration^29^, we reduced the HARBOR AAV dose to one-tenth of that used for 3D5. Despite the substantially lower dose, HARBOR achieved even higher *circPOLR2A* expression levels in the hippocampus (Extended Data Fig. 3c). Consistent with cellular results (Extended Data Fig. 3b), both HARBOR-*circPOLR2A(9,10)* and 3D5-*circPOLR2A(9,10)* effectively repressed hippocampal PKR activation and pro-inflammatory cytokine expression relative to EGFP controls in 5×FAD AD mice (Extended Data Fig. 3d, 3e). Histopathological analyses revealed reduced Aβ plaque accumulation (Fig. 4e, 4f; Extended Data Fig. 3d), attenuated microglial and astrocytic overactivation (Fig. 4e, 4f; Extended Data Fig. 3f, 3g), and preserved neuronal integrity (Fig. 4e, 4f) in both treatment groups. Notably, HARBOR achieved better neuroprotective effects than 3D5 (Fig. 4e, 4f; Extended Data Fig. 3f, 3g), despite the 10-fold lower viral dosage (Extended Data Fig. 3c).

We also examined the therapeutic efficacy of the HARBOR-derived *circPOLR2A(9,10)* in 8-month-old PS19 mice, another neurodegenerative disease model characterized primarily by Tau hyperphosphorylation^30^. Mice were injected with AAV-EGFP as a control, or *circPOLR2A(9,10)* expressed by HARBOR or 3D5 systems, with HARBOR administered at one-tenth the viral dose used for 3D5 (Extended Data Fig. 4a). Both HARBOR- and 3D5-derived *circPOLR2A(9,10)* significantly reduced glial activation in PS19 mice compared with the EGFP group (Extended Data Fig. 4b, 4c), accompanied by decreased hippocampal Tau hyperphosphorylation (Extended Data Fig. 4c, 4d) and improved neuronal preservation (Extended Data Fig. 4e, 4f). with HARBOR achieving greater effects despite the substantially lower viral dose, supporting HARBOR as a simple and robust circRNA aptamer expression platform with superior therapeutic efficacy *in vivo*.

### HARBOR-derived circular RNA aptamers inhibit pathological TDP-43 condensation

Pathological TDP-43 condensation and aggregation is a hallmark of multiple neurodegenerative disorders including amyotrophic lateral sclerosis (ALS) and frontotemporal dementia (FTD)^31, 32^. Our recent study has shown that the loosely structured *circT3* (332 nt core sequence) and *circT3-M3* aptamers (89 nt core sequence) with 102 nt and 105 nt single-stranded region, respectively, can efficiently prevent pathological TDP-43 condensation in cytosol^33^. To further evaluate the versatile application of HARBOR as an efficient platform to deliver therapeutic circRNA aptamers, we generated HARBOR constructs encoding *circT3* and *circT3-M3* and evaluated their capacity in suppressing TDP-43 condensation in cell (Fig. 4g, 4h).

To achieve this, we employed a well-established cellular model in which U2OS cells were transfected with mNG-tagged NLS-mutant TDP-43 (mNG-TDP-43^NLS^) and subsequently treated with 250 μM sodium arsenite (SA) for 1 h to induce cytosolic TDP-43 condensation formation^33–35^. Under these conditions, U2OS cells expressing mNG-TDP-43^NLS^ alone and treated with SA exhibit robust cytosolic TDP-43 condensation in 94% of cells, consistent with the previously reported model^33^ (Fig. 4i, 4j). For functional assessment, U2OS cells were co-transfected with mNG-TDP-43^NLS^ together with HARBOR-expressed *circT3*, *circT3-M3*, or a non-targeting *T9* control RNA^33^, followed by SA treatment under the same conditions (Fig. 4h). Imaging analysis showed that *circT3* and *circT3-M3* expressed by HARBOR markedly inhibited TDP-43 condensation with less than 10% of the cells displayed cytosolic TDP-43 condensation (Fig. 4i, 4j). As a negative control, the HARBOR-expressed non-targeting T9 RNA^33^ failed to inhibit TDP-43 condensation (Fig. 4i, 4j). Together, these results demonstrate HARBOR-driven aptamer expression efficiently suppresses pathological TDP-43 condensation, supporting its potential for TDP-43 proteinopathy therapeutics.

### HARBOR enables efficient base editing

A major challenge of endogenous circRNA expression systems is their poor efficiency in producing large, translatable circRNAs^6, 18^. This constraint has restricted their application in genome editing, as most base editors consist of large Cas-derived proteins exceeding several kilobases in coding length^36, 37^. Given the markedly enhanced circularization efficiency of cargo and robust protein express of HARBOR (Fig. 3), we evaluated its suitability for expressing functional base editors.

We generated HARBOR-derived circRNA constructs encoding *CVB3-ABE8e*, a 5,553-nt circRNA cargo expressing ABE8e, an evolved adenine base editor consisting of a hyperactive adenine deaminase fused to SpCas9 nickase^36^, designed for A•T-to-G•C base conversion (Fig. 5a). Immunoblotting confirmed efficient ABE8e protein production from HARBOR (Fig. 5b). To assess editing activity, we selected several well-established benchmark target sites (site2, site3, and RNF2) that have been extensively validated in previous studies for quantifying base-editing efficiency^37^. By co-expression of HARBOR-ABE8e with corresponding guide RNAs in 293T cells, we found that all of them displayed efficient base editing at their target sites with the efficiency of ∼35%-70% (Fig. 5c). Next, we applied HARBOR to express saKKH-ABE8e, a 5,553 nt large circRNA cargo expressing a compact *Staphylococcus aureus* Cas9-based ABE variant with expanded PAM compatibility^36^. We found that HARBOR-driven saKKH-ABE8e also yielded efficient protein production and efficient base editing, achieving ∼40-60% editing at the previously validated benchmark loci site9 and site11^36^ (Fig. 5d, 5e). These results demonstrate that HARBOR can effectively support both the expression and functional activity of large genome-editing effectors.

**Fig. 5.**
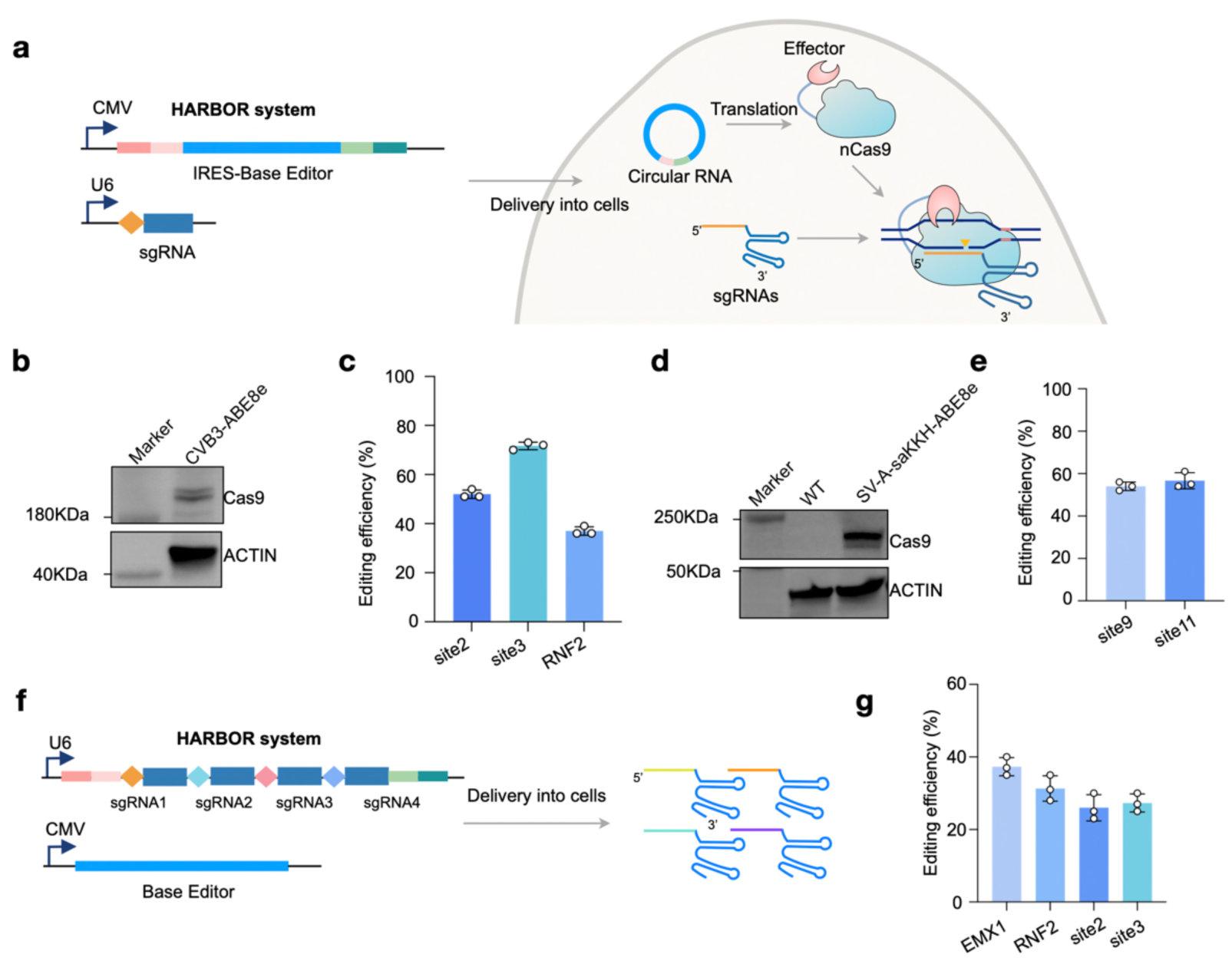
HARBOR enables functional protein expression for base editing. **a**, Schematic of HARBOR-mediated expression of Base Editors (ABE8e and saKKH-ABE8e^36^) for genome editing. **b-c**, HARBOR enables an efficient expression of the adenine base editor ABE8e. **b**, Western blotting confirms ABE8e protein expression in 293T cells. **c**, Editing efficiencies of ABE8e at the site2, site3 and RNF2 target loci. **d-e**, HARBOR enables an efficient expression of the base editor saKKH-ABE8e. **d**, Western blotting shows protein expression of saKKH-ABE8e in 293T cells. **e**, Editing efficiencies of saKKH-ABE8e at two genomic loci, site9 and site11. **f**-**g**, HARBOR enables efficient expression of gRNA arrays. **f**, Schematic of gRNA arrays for genome editing expressed by HARBOR. **g**, Editing efficiencies of gRNA arrays at four genomic loci, including EMX1, RNF2, site2, and site3. In panel **c**, **e** and **g**, error bars represent the mean ± s.d.

Multiplexed guide RNA expression is critical for many genome-engineering applications, including simultaneous editing of multiple loci, combinatorial genetic perturbation, and therapeutic correction of complex genotypes^38, 39^. Given HAROR can generate large-sized circRNAs (Fig.3), we also evaluated its capacity for such a purpose. We constructed a tandem array encoding four sgRNAs targeting EMX1, RNF2, site2, and site3, all of which represent widely used benchmark loci for evaluating base editing performance^37^ (Fig. 5f). Sequencing confirmed efficient editing across all four targets, with editing efficiencies of ∼25%-40% (Fig. 5g). Thus, HARBOR effectively supports multi-guide RNA arrays, achieving editing efficiencies comparable to those obtained via separate expression of editing effectors and gRNAs (Fig. 5a–5e). Together, these results demonstrate HARBOR’s versatility for expressing large editing effectors and supporting complex multiplexed genome engineering.

### HARBOR enables fibroblast trans-differentiation

To further expand the application potential of HARBOR, we assessed its capability to drive transcription factor-mediated cell fate conversion. Myod1, a master regulator of myogenesis known to convert fibroblasts into myogenic lineage cells^40, 41^, was expressed from HARBOR-derived circRNAs in mouse 10T1/2 fibroblasts (Fig. 6a). Following transfection, cells were cultured in differentiation medium, resulting in time-dependent induction of myogenic marker genes and proteins (Fig. 6b, 6c). Immunofluorescence confirmed the formation of multinucleated myotube-like structures positive for myogenic differentiation marker α-Actinin at day 7 (Fig. 6d), showing that HARBOR-derived circRNA translation is sufficient to drive functional cellular trans-differentiation.

**Fig. 6.**
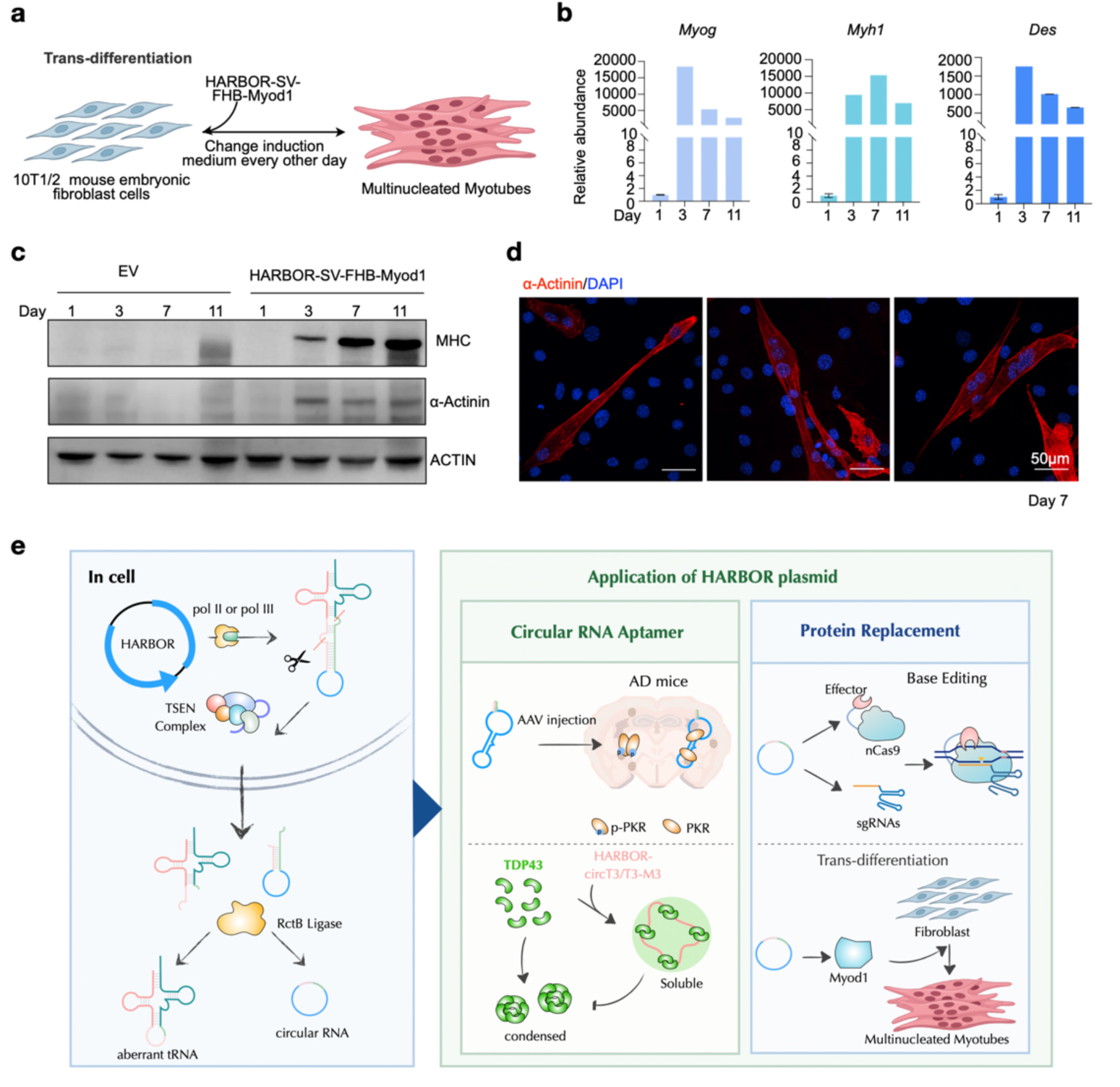
HARBOR enables functional protein expression for trans-differentiation. **a**, Schematic of fibroblast cells trans-differentiation into multinucleated myotubes following Myod1^40, 41, 47^ expression by HARBOR vector. **b**, qPCR analysis of myogenic marker gene expression (Myog, Myh1, and Des) at day 1, 3, 7, and 11 following Myod1-induced trans-differentiation mediated by HARBOR. **c**. Western blotting confirms the expression of myogenic marker proteins MHC and α-Actinin following trans-differentiation at day 1, 3, 7, and 11. **d**, Immunofluorescence of α-Actinin expression and cellular morphology at day 1, 3, 7, and 11 of fibroblast cells trans-differentiation. α-Actinin is shown in red, and nuclei are stained with DAPI (blue). Scale bar, 50 μm. **e**, Summary of HARBOR-mediated circRNA biogenesis and its applications for functional RNA aptamers and protein replacement. In panel **b**, error bars represent the mean ± s.d.

In sum, HARBOR expands the design landscape of circRNA engineering via a modular, structurally defined expression architecture that unifies high-efficiency circularization of noncoding and coding RNA cargos (Fig. 6e). This robust platform will advance a broader application of circRNA technologies in modulating gene regulation, cellular engineering, and RNA-based therapeutics.

## Discussion

An efficient and robust platform for functional circRNA expression comprising both the aptamer and translating modes in one platform in cells has been lacking. Here, we establish HARBOR, a circRNA expression platform built upon a human pre-tRNA scaffold integrated with an archaeal BHB motif, enabling robust and efficient RNA circularization in cells (Fig.1, 2) and *in vivo* (Fig. 4). Compared with previously reported back-splicing-^11, 12, 14^, tRNA intron splicing-^16, 17^, and ribozyme-mediated circRNA expression systems^6, 18^, HARBOR supports efficient production of therapeutic aptamers (Fig.4), circularization of RNA sequences up to 8 knt (Fig. 3a, 3b and Extended Data Fig. 2a) and drives markedly elevated protein in cells (Fig. 3i–3l). The combination of broad cargo compatibility and enhanced translational capacity provides a generalizable framework for circRNA engineering, thus extending the functional applicability of circRNA-based modalities (Fig. 4, 5 and 6).

A key advantage that distinguishes the HARBOR platform from existing circRNA expression strategies lies in its structure-guided mode of RNA processing. Canonical back-splicing systems rely on splice-site recognition and often exhibit strong sequence-context dependency^11–14^, whereas ribozyme-based approaches require precise folding and catalytic activity of the ribozyme itself^6, 18^. In contrast, HARBOR utilizes endogenous tRNA splicing machinery to process a defined BHB structure^42, 43^, eliminating the requirement for canonical exon-intron organization and reducing the need for extensive sequence optimization. Moreover, the extensive duplex introduced at tRNA intron would not interfere with BHB cleavage as this region is not recognized by human TSEN^24–26^, but can enhance the folding and BHB motif formation of cargo-loaded pre-tRNA and the circularization efficiency of cleaved intron. We speculate that this structural designing principle contributes to the robust circularization observed across diverse cargo sequences and facilitates efficient processing of a variety of RNA transcripts. These findings expand the design principles available for circRNA engineering and establish BHB-directed circularization as a broadly applicable approach for intracellular circRNA production.

Beyond improving circRNA expression capacity, HARBOR also expands the functional scope of engineered circRNAs. We demonstrate its utility in both non-coding and coding applications, including aptamer-targeted alleviation of pathological proteins in different contexts (Fig. 4), efficient adenine base editing through circRNA-encoded ABE8e and saKKH-ABE8e (Fig. 5), and fibroblast-to-myocyte trans-differentiation driven by circRNA-expressed MYOD1 (Fig. 6). These examples highlight the versatility of HARBOR-derived circRNAs as durable intracellular functional modules. Given their enhanced stability and sustained activity, HARBOR-generated circRNAs are expected to be adapted for a wide range of therapeutic applications, including protein replacement, programmable genome engineering, cell reprogramming, molecular decoys, RNA scaffolds, and synthetic gene regulatory circuits.

Despite these advantages, the current HARBOR platform has some limitations. While it exhibits reliable performance in cellular and animal models, systematic *in vivo* characterization is required to profile its tissue distribution, and overall biocompatibility. Although no overt cytotoxicity was detected throughout our experimental conditions, engineered circRNAs harboring functional binding sites for RNA-binding proteins or microRNAs^44–46^ may engage endogenous cellular regulatory networks, potentially inducing context-dependent off-target effects. Careful evaluation of functional specificity of HARBOR-derived circRNAs under future applications is expected for translational development and therapeutic deployment.

## Methods and Materials

### Plasmid constructions

To construct vectors for endogenous circRNA overexpression, different versions of the BHB motif and pre-tRNA scaffolds carrying the sequence of interest were cloned downstream of either the U6 or CMV promoter in a pAAV-based or p23-based backbone plasmid. For the 3D5 and Tornado systems, cargo sequences were inserted into the corresponding backbone vectors by homologous recombination. All constructs were verified by Sanger sequencing. Detailed sequence information for all constructs is provided in Supplementary Table 1.

### Experimental animals

5×FAD mice on a C57BL/6J background were generously provided by NaiHe Jing at the Guangzhou Institute of Biomedicine and Health, Chinese Academy of Sciences (CAS). PS19 mice on a C57BL/6J background were generously provided by Yuejun Chen and Shiqing Cai at the Center for Excellence in Brain Science and Intelligence Technology, CAS. All animal procedures were approved by the Institutional Animal Care and Use Committee of the Center for Excellence in Molecular Cell Science (CEMCS), CAS.

For AAV-mediated delivery experiments, AD mice received intracerebral AAV injections at 8 months of age. Molecular and histological analyses were performed 1 month after injection.

### Virus strains

The overexpressed virus AAV-PHP.eB was packaged and produced by Guangzhou PackGene Biotech Co., Ltd.

### Cell

HeLa and HEK293FT cells were purchased from the American Type Culture Collection (ATCC; http://www.atcc.org). A549, SH-SY5Y, HepG2, HCT116, HeLa, C3H/10T1/2, 293T and U2OS cells were purchased from National Collection of Authenticated Cell Cultures. All cell lines were authenticated by short tandem repeat (STR) profiling.

HeLa, C3H/10T1/2 and HEK293FT cells were cultured in DMEM supplemented with 10% fetal bovine serum (FBS). A549 cells were maintained in F-12K medium supplemented with 10% FBS and 0.1 mM GlutaMax. SH-SY5Y cells were maintained in a 1:1 mixture of MEM and F12 medium supplemented with 10% FBS, 0.1 mM L-Glutamax, 0.1 mM non-essential amino acids (NEAA) and 1 mM Sodium pyruvate. HCT116 cells were maintained in RPMI-1640 supplemented with 10% FBS. HepG2 cells were maintained in MEM supplemented with 10% FBS and 0.1 mM NEAA. U2OS cells were maintained in McCoy’s 5a medium supplemented with 10% FBS. All cell lines were maintained at 37°C in a 5% CO_2_ cell culture incubator and routinely tested tested for mycoplasma contamination.

### Cell transfection and lentivirus infection

For plasmid transfection, cells were seeded into 6-well plates 1 day before transfection. When cell confluency reached approximately 50% the following day, plasmids were transfected using Lipofectamine 3000 (Invitrogen) according to the manufacturer’s instructions.

For lentiviral infection, C3H/10T1/2 cells were dissociated into single cells using Trypsin-EDTA (Gibco) and cultured in the corresponding growth medium supplemented with 10 μL concentrated lentivirus and 10 μg ml⁻¹ polybrene. To enhance infection efficiency, cells were cultured for an additional day in the presence of 1 μg ml⁻¹ puromycin starting 2 days after infection.

### Lentivirus production

To produce lentiviral particles, 5 × 10^6^ C3H/10T1/2 cells in a 10 cm dish were co-transfected with 10 μg pLKO.1-shRNA, 7.5 μg psPAX2 and 3 μg pMD2.G plasmids. The supernatant containing viral particles was harvested at 48 hrs and 72 hrs after transfection, then filtered through Millex-GP Filter Unit (0.22 μm pore size, Millipore). Viral particles were concentrated 100-fold by Lenti-Concention Virus Precipitation Solution (ExCell Bio), resuspended in PBS containing 0.1% BSA, and stored at -80°C until use.

### RNA isolation, RT-qPCR

Total RNAs from cultured cells or cultured cells with different treatments were extracted with Trizol Reagent (Invitrogen) according to the manufacturer’s protocol. The reverse transcription was carried out using PrimeScript RT Master Mix (TaKaRa) according to the manufacturer’s protocol. RT-qPCR was performed using SYBR Green Realtime PCR Master Mix (TOYOBO) and the StepOnePlus real-time PCR system (Applied Biosysterms). The relative expression of different sets of genes was quantified to *GAPDH* mRNA. All primers used for RT-qPCR were listed in Supplementary Table 1.

### Northern blotting (NB)

NB was carried out according to the manufacturer’s protocol (DIG Northern Starter Kit, Roche). Briefly, 5 μg of total RNA was loaded on 1% agarose gel or denatured PAGE gel. Electrophoresis was performed at 120 V for 120 min at 4℃. RNA was then transferred onto a nylon membrane (Roche) under constant 20 V for 20 hrs at 4℃ or 400 mA for 90 min at 4℃. Following transfer, RNA was ultraviolet (UV) cross-linked and prehybridized with 3 mL DIG Easy Hybridization Granules (Roche) at 68°C for 3 hrs. Digoxigenin (Dig) labeled RNA probes were generated using T7 RNA polymerase by *in vitro* transcription (IVT) with the RiboMAX Large Scale RNA Production System (Promega). RNA probes were denatured at 95℃ for 5 min and cooled on ice for 2 min. For RNA hybridization, 800ng of the corresponding RNA probe was diluted into 3 mL DIG Easy Hybridization Granules and incubated with the membrane at 68°C for 16 hrs. The next day, the membrane was washed twice at RT with 2 × SSC containing 0.1% SDS for 5 min followed by two washes at 68℃ with 0.2 × SSC containing 0.1% SDS for 30 min. The membrane was then washed at RT for 5 min with washing buffer containing 0.3% Tween 20 and 1 × maleic acid. The membrane was blocked with 1 × blocking buffer for 30 min. Anti-DIG antibody was prepared by diluting Anti-DIG-AP (Roche) into the 1 × blocking buffer. After blocking, the membrane was incubated with the diluted Anti-DIG-AP at RT for 30 min followed by three 20-minute washes with washing buffer and incubation with 1X detection buffer. Chemiluminescence imaging was done using CDP-Star (Roche) and captured using the Sagecreation imaging system. All primers used for making probes were listed in Supplementary Table 1.

### Immunofluorescence (IF)

Cells were fixed in 4% paraformaldehyde (PFA) for 10 min followed by 3 washes with 1× PBS. The cells were then permeabilized in 0.5% Triton X-100 in 1× PBS for 10 min. After blocking with 1% BSA for 30 min, cells were incubated with specific primary antibodies anti-α-Actinin (1:100, Abcam) overnight at 4°C. The next day, cells were washed 3 times with 1× PBS for 5 min followed by the incubation of secondary antibody for 1 hr at RT in the dark. Nuclei were counterstained with DAPI for 2 min at RT in the dark followed by 3 washes with 1× PBS. 10 μL of VECTASHIELD® PLUS Antifade Mount Medium (Vector) was added onto the slide, followed by the cover slip. The coverslip was sealed to the slide with nail polish to prevent evaporation and introduction of oxygen. The images were collected on a Nikon spatial array confocal microscope.

Floating brains sectioned and stored in PBS were blocked with 10% donkey serum, 0.125% gelatin and 0.05% Tween-20 in PBS for 2 h at room temperature before applying the primary antibody master mix diluted in antibody diluent solution (1% BSA and 0.03% Triton X-100 in PBS) overnight at 4 °C. Samples were stained with anti-Aβ (H31L21, Invitrogen, 1:1,000; 6Ε10, BioLegend, 1:1,000) to label plaques. To study microglial and astrocyte morphology and density, as well as neuronal numbers, sections were stained with Iba1 (1:1,000, Wako), GFAP (1:1,000, CST) and NeuN (1:1000, Abcam). To assess tau pathology in PS19 mice, brain sections were immunostained with anti-phospho-tau (1:500, Thermo Fisher). Sections were then washed three times for 10 min at room temperature in PBS and then incubated in matched donkey anti-rabbit, anti-goat and anti-mouse labeled with Alexa Fluor 488, 594 and 647 (Jackson, 1:1,000 dilution) at 37 °C for 30 min. Samples were washed again three times for 10 min at room temperature and incubated with DAPI (1:1,000) for 10 min at room temperature All tissue sections were then transferred to wells containing PBS before mounting to glass slides with VECTASHIELD antifade mounting medium (Vector Lab) and coverslips. Mounted slides were stored at 4 °C until imaging using Nikon spatial array confocal microscope. Quantification of images was performed using Fiji software.

### Tissue processing and sample preparation

For immunofluorescence, mice were euthanized and transcardially perfused with 1× PBS (25 ml) followed by 4% PFA (25 ml at 37 °C and 25 ml at 4 °C). Brains were dissected and post-fixed in 4% PFA overnight at 4 °C, then transferred to 25% sucrose for 24 h and 30% sucrose for 48 h. Tissues were embedded in Epredia Neg-50 (Thermo Fisher Scientific), frozen, and sectioned coronally at 25 μm using a Leica cryostat. Free-floating sections were stored in PBS at 4 °C for subsequent staining.

For protein analyses, hippocampus and cortex were rapidly dissected, snap-frozen, and stored at −80 °C. Hippocampal tissues were homogenized in 300 μl RIPA buffer (Beyotime) with protease and phosphatase inhibitor cocktail (Beyotime). Lysates were centrifuged at 15,000 rpm for 10 min, and supernatants were collected for Western blot. For RNA extraction, tissues were homogenized directly in TRIzol using a tissue grinder. After centrifugation at 15,000 rpm for 10 min, the aqueous phase was separated, and chloroform was added to proceed with standard phase separation and subsequent RNA isolation.

### Western blotting (WB)

Cell lysates or tissues were mixed with 1× SDS loading buffer (50 mM Tris-HCl, pH 6.8, 2% SDS, 10% glycerol, 0.05% bromophenol blue, 5% 2-Mercaptoethanol). Samples were boiled at 95℃ in a metal bath for 10 min, followed by ice cooling for 2 min. After quick centrifugation, the samples were loaded on the SDS-PAGE gel for electrophoresis and transferred onto a PVDF membrane. Following transfer, the membrane was incubated with primary antibody overnight at 4℃, washed 3 times with PBST for 10 min, incubated with secondary antibody for 1 hr at RT, washed 3 times with PBST for 10 min, then incubated with the ECL detection buffers (Thermo Fisher Scientific). The protein signals were detected via eBlot Touch Imager and the intensity of the bands was quantified using Fiji software.

Specifically, for phosphorylated protein detection, cell or tissue was lysed in RIPA lysis buffer supplemented with protease and phosphatase inhibitor cocktail. For cells, the sample was centrifuged, and the supernatant was collected, mixed with 5× loading buffer and heated at 95 °C. For tissue samples, after 90 s of homogenization, the sample was centrifuged, and the supernatant was collected. The samples were then mixed with 5× loading buffer and heated at 95 °C.

Primary antibodies used were anti-FLAG (1:1,000, Sigma), anti-Myosin heavy chain (1:1000 ABclonal), anti-α-Actinin (1:1000, Abcam), anti-Cas9 (1:1000, CST), anti-PKR (1:1000, Abways), anti-phospho-PKR (1:1000, Abways), anti-PKR (1:200, Santa Cruz), anti phospho-PKR (1:1000, Sigma), anti-β-Amyloid (1:1000, Biolegend), anti-Aβ plaque (1:1000, Biolegend) and anti-β-actin (1:1,000, Sigma).

### Base editing analysis

Genomic DNA was extracted from cultured cells using the TIANamp Genomic DNA Kit (TIANGEN). Target loci were amplified by PCR using locus-specific primers and subjected to Sanger sequencing for editing analysis. Base-editing efficiency was quantified using EditR analysis by calculating the percentage of edited nucleotides at the target site.

### MyoD1-induced transdifferentiation of C3H/10T1/2 cells

C3H/10T1/2 cells were seeded 1 day before infection and cultured in growth medium until reaching approximately 50-70% confluency. Cells were then transduced with lentivirus expressing MyoD1 in the presence of 10 μg ml⁻¹ polybrene. Twenty-four hours after infection, the medium was replaced with fresh growth medium, and infected cells were selected with 1 μg ml⁻¹ puromycin beginning 48 h post-infection.

To induce myogenic transdifferentiation, cells were switched to differentiation medium consisting of DMEM supplemented with 2% horse serum when reaching near confluency. The differentiation medium was replaced every 2 days. Myogenic differentiation was assessed by morphological analysis, RT-qPCR, western blot and immunostaining for myogenic markers at the indicated time points.

#### Acknowledgements

We thank N. Jing for gifting 5×FAD mice and S. Cai and Y. Chen for gifting PS19 mice. S. R. Jaffrey for gifting Tornado vector. This work was supported by the National Key R&D Program of China (2021YFA1300501), Strategic Priority Research Program of the CAS (XDB0570000) and Science and Technology Commission of Shanghai Municipality (23DX1900100 and 23DX1900101) to L.-L.C. China National Postdoctoral Fellowship Program (Grade B) (GZB20250592), China Postdoctoral Science Foundation (2025M782805) and Shanghai Post-doctoral Excellence Program (2025194) to S.-M.C. This work has been supported by the New Cornerstone Science Foundation through the New Cornerstone Investigator Program and the Ruisi Research Center for Life Science, Minhang District, Shanghai. L.-L.C. is also a Shanghai Academy of Natural Sciences Senior Investigator. The Author Xin Feng Gratefully Acknowledges the Support of Sanofi Scholarship Program.

## Author contributions

L.-L.C. and K.Y. conceptualized the project. L.-L.C. supervised the project. S.-M.C., X.F., J.T.-C.Z., H.W. performed the experiments. X.W. provided plasmids related to base editing. S.-M.C., X.F., J.T.-C.Z. and H.W. wrote the paper with input from all authors. L.-L.C. and K.Y. edited the manuscript. All authors read and approved the manuscript.

## Extended Data and Extended Data Figure Legends

**Extended Data Fig. 1.**
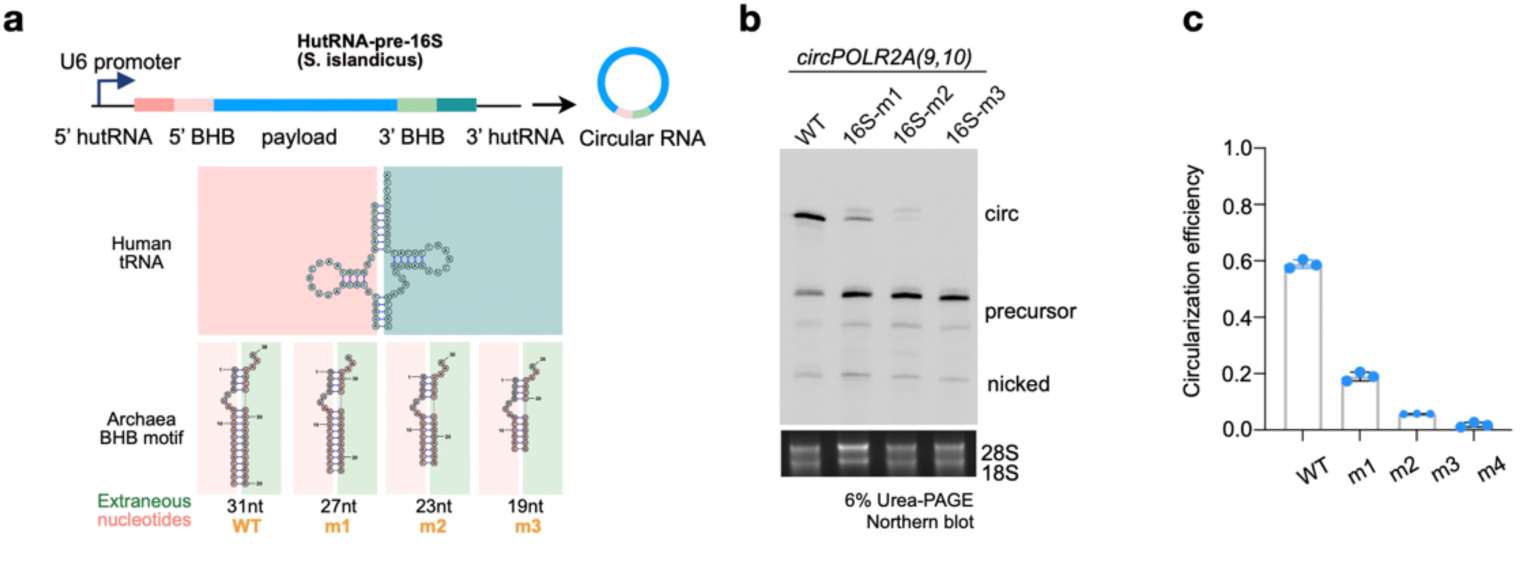
Truncation of hutRNA-pre-16S reduces circularization efficiency. **a**, Schematic illustration of truncation mutations in the hutRNA-pre16S plasmid construct, in which the long double-stranded RNA paring region is progressively shortened by four nucleotides and designated as m1, m2, and m3 respectively. **b**, Northern blotting analysis of circular RNA formation mediated by different truncation mutants shown in panel **a**. “circ” is RNA circle; “precursor” is linear precursor; “nicked” is nicked circle. **c**, Quantification of circular RNA circularization efficiency mediated by different plasmid vectors shown in panel **b**. Error bars represent the mean ± s.d.

**Extended Data Fig. 2.**
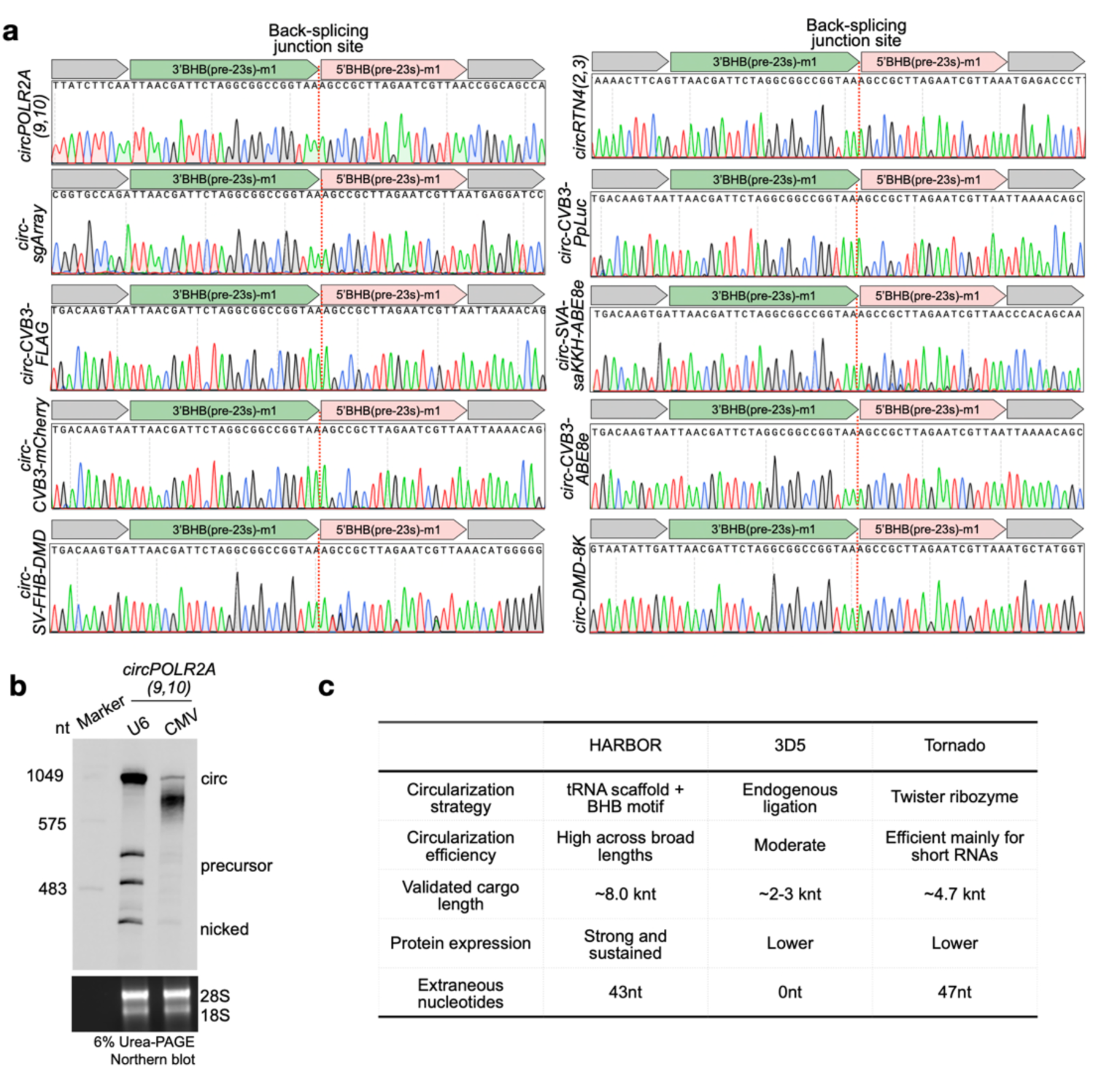
Comparison of HARBOR, 3D5, and Tornado circular RNA expression systems. **a**, Sanger sequencing confirmed the back-splice junction of eight circular RNAs of different lengths (336-8,000 nt) generated by HARBOR, including *circPOLR2A(9,10)*, *circ-sgArray*, *circ-CVB3-FLAG*, *circ-CVB3-mCherry*, *circRTN4(2,3)*, *circ-CVB3-PpLuc*, *circ-SV-A-saKKH-ABE8e*, *circ-CVB3-ABE8e*, *circ-SV-FHB-DMD* and *circ-DMD-8K*. **b**, CMV promoter-driven expression yields low circularization efficiency of short RNA-*circPOLR2A(9,10)* constructs. Cells were transfected with U6- or CMV-driven RNA-*circPOLR2A(9,10)* constructs. Total RNA was analyzed by Northern blotting to assess circular RNA formation. CMV-driven expression shows lower circularization efficiency compared to U6. **c**, Comparison of performance features among three endogenous circularization vectors: HARBOR, 3D5, and Tornado.

**Extended Data Fig. 3.**
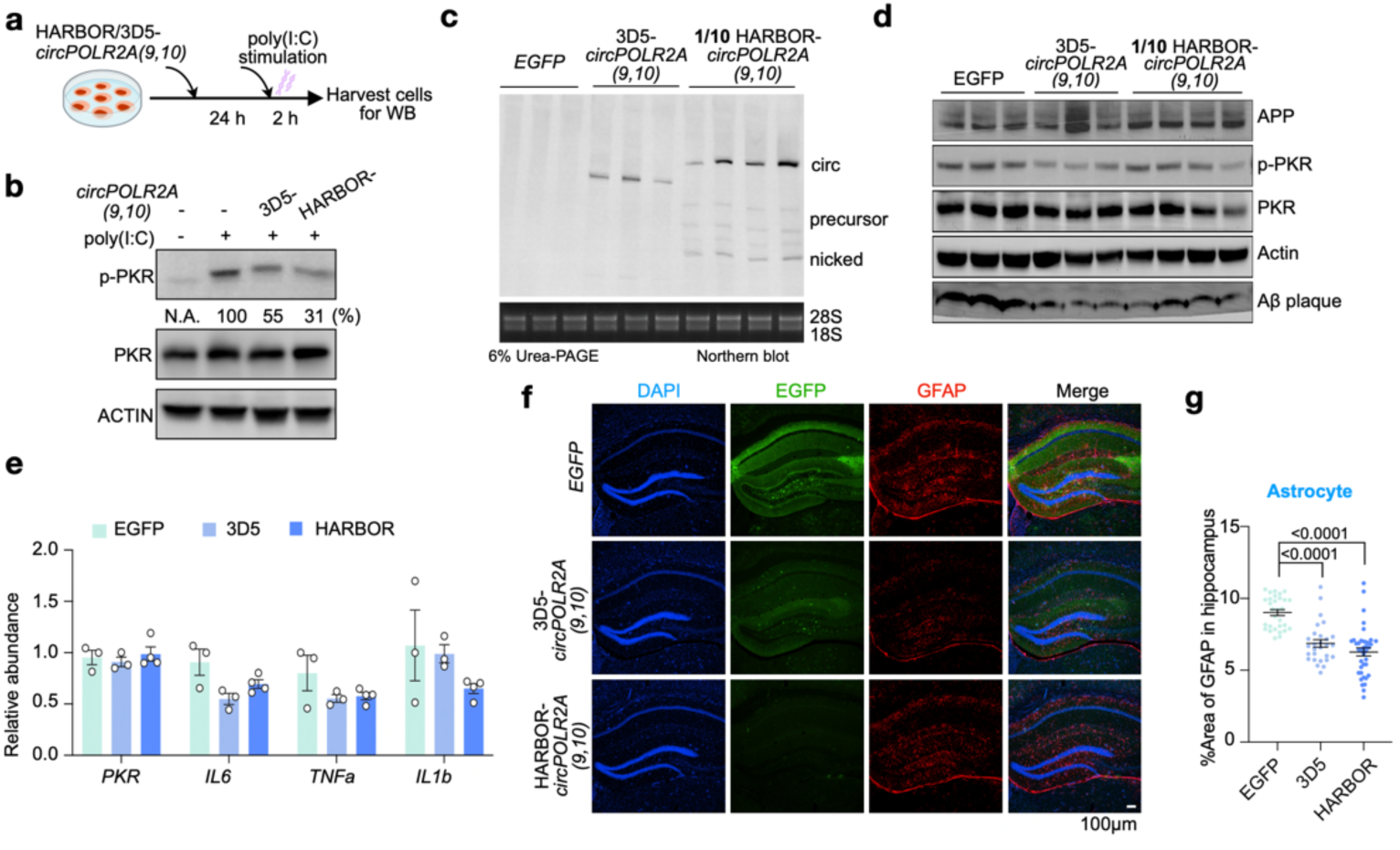
HARBOR-expressed ds-cRNAs alleviate neuroinflammation in 5×FAD mice. **a**-**b**, HARBOR-derived *circPOLR2A(9,10)* suppresses PKR phosphorylation in cells more effectively than its 3D5-expressed counterpart. **a**, A549 cells were individually transfected with 3D5- or HARBOR-*circPOLR2A(9,10)* and cultured for 24 h, followed by poly(I:C) stimulation. **b**, PKR phosphorylation (p-PKR at T446; p-PKR for simplicity) and total PKR expression were assessed. The levels of p-PKR in each panel were quantified using image J, and p-PKR levels were normalized to total PKR, respectively. **c**, AAV-HARBOR drives much higher circRNA expression than AAV-3D5. A 10-fold diluted AAV-HARBOR achieved circRNA levels comparable to AAV-3D5 in hippocampus, shown by Northern blotting. **d**, Western blotting analysis showing that HARBOR-*circPOLR2A(9,10)* at one-tenth the dosage achieves suppression of excessive PKR phosphorylation and reduces APP and Aβ plaque levels in the hippocampus of 5×FAD mice comparable to that of 3D5-*circPOLR2A(9,10)*. **e**, Expression of proinflammatory cytokine genes (*Il6*, *Tnfa* and *Il1b*) are reduced in both the 3D5 and HARBOR groups compared with the EGFP control in the hippocampus of 5×FAD mice one month after AAV injection. RT-qPCR analysis was performed to quantify *Pkr* and cytokine transcript levels in EGFP, 3D5- and HARBOR-*circPOLR2A(9,10)* treated hippocampal tissues. n = 3 biologically independent mice per group. **f**-**g**, Alleviated gliosis in the hippocampus of 5**×**FAD mice 1 month after AAV-mediated delivery of HARBOR-*circPOLR2A(9,10)* or 3D5-*circPOLR2A(9,10)*, with HARBOR-*circPOLR2A(9,10)* showing a stronger effect. **f**, Representative images of hippocampal sections from 5×FAD mice injected AAV-EGFP, AAV-3D5- or HARBOR-*circPOLR2A(9,10)*, stained with DAPI (blue), EGFP (green), and GFAP (red). Scale bar, 100 μm. **g**, Quantification showing reduced astrocyte activation relative to the EGFP control group, with a greater effect observed in the AAV-HARBOR-*circPOLR2A(9,10)* group than in the AAV-3D5-*circPOLR2A(9,10)* group. In panel **e** and **g**, statistical analysis was performed using a two-tailed Student’s t-test. Error bars represent the mean ± s.e.m.

**Extended Data Fig. 4.**
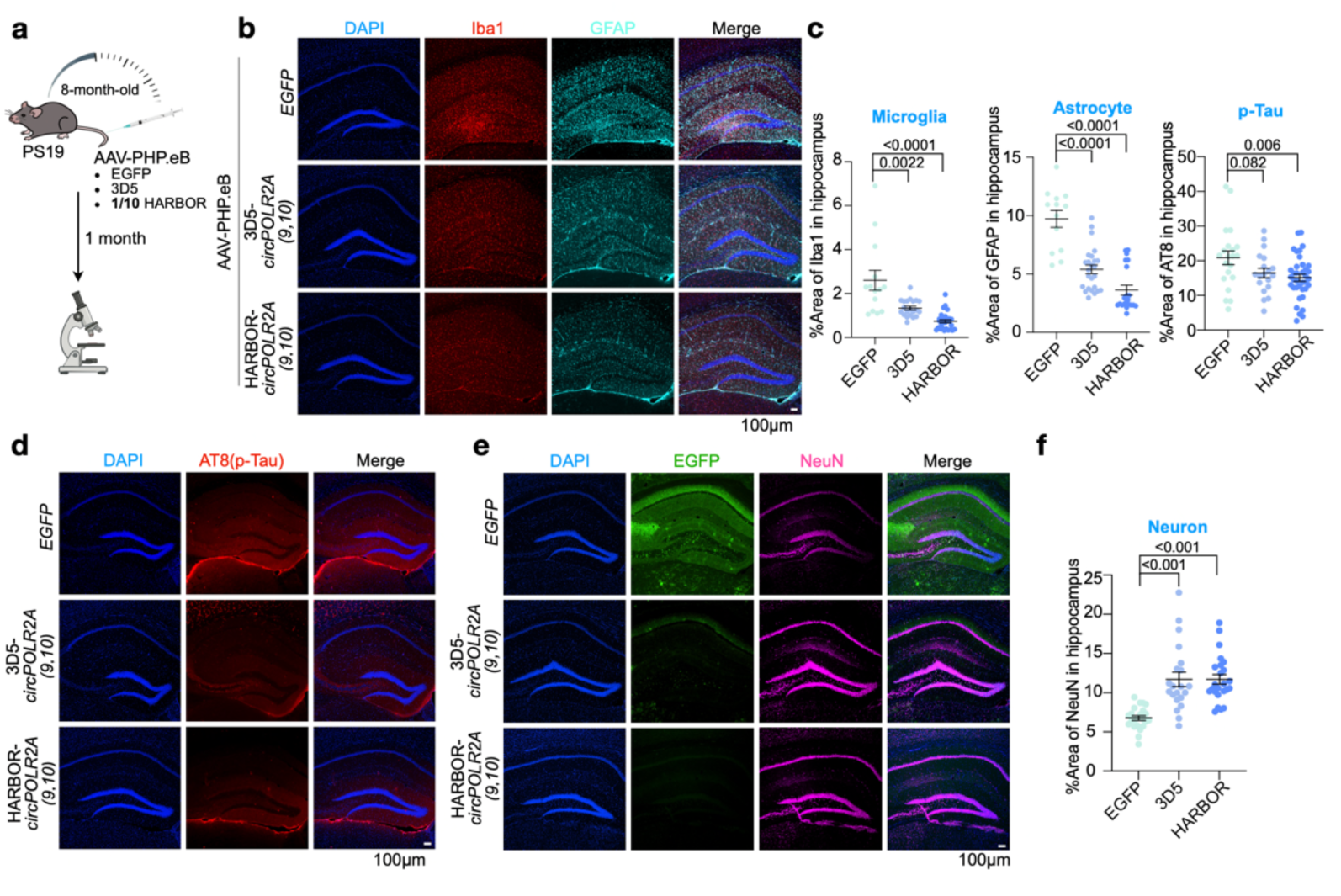
HARBOR-expressed ds-cRNAs alleviate neuroinflammation in PS19 mice. **a**, Schematic illustration of the experimental design. AAV-PHP.eB-delivered HARBOR-*circPOLR2A(9,10)* was injected into the hippocampus of PS19 mice. Samples were collected one month after AAV injection for analysis. **b**-**d**, Alleviated gliosis and reduced Tau phosphorylation in the hippocampus of PS19 mice 1 month after AAV-mediated delivery of HARBOR-*circPOLR2A(9,10)* or AAV-3D5-*circPOLR2A(9,10)*, with HARBOR-*circPOLR2A(9,10)* showing a stronger effect. **b**, Representative images of hippocampal sections from PS19 mice one month after AAV injection with AAV-EGFP, AAV-3D5- or HARBOR-*circPOLR2A(9,10)*, stained with DAPI (blue), Iba1 (red), and GFAP (cyan). Scale bar, 100 μm. **c**, Quantification showing reduced microglia and astrocyte activation relative to the EGFP control group, with a greater effect observed in the AAV-HARBOR-*circPOLR2A(9,10)* group than in the AAV-3D5-*circPOLR2A(9,10)* group. **d**, Representative images of hippocampal sections from PS19 mice injected AAV-EGFP, AAV-3D5- or HARBOR-*circPOLR2A(9,10)*, stained with DAPI (blue) and Tau phosphorylation (red, AT8). Scale bar, 100 μm. **e**-**f**, Reduced neuronal loss in the hippocampus of PS19 mice 1 month after AAV-delivered HARBOR-*circPOLR2A(9,10)* or AAV-3D5-*circPOLR2A(9,10)* injection, with HARBOR-*circPOLR2A(9,10)* showing a stronger effect. **e**, Representative images of hippocampal sections from PS19 mice injected AAV-EGFP, AAV-3D5- or HARBOR-*circPOLR2A(9,10)*, stained with DAPI (blue), EGFP (green) and NeuN (magenta). Scale bar, 100 μm. **f**, Quantification reveals reduced neuronal loss in the AAV-HARBOR- and AAV-3D5-*circPOLR2A(9,10)* groups compared with the EGFP control. In panel **c** and **f**, statistical analysis was performed using a two-tailed Student’s t-test. Error bars represent the mean ± s.e.m.

